# Finemap-MiXeR: A variational Bayesian approach for genetic finemapping

**DOI:** 10.1101/2022.11.30.518509

**Authors:** Bayram Cevdet Akdeniz, Oleksandr Frei, Alexey Shadrin, Dmitry Vetrov, Dmitry Kropotov, Eivind Hovig, Ole A. Andreassen, Anders M. Dale

**Affiliations:** NORMENT Centre, Division of Mental Health and Addiction, Oslo University Hospital & Institute of Clinical Medicine, University of Oslo, Oslo, Norway; Centre for Bioinformatics, Department of Informatics, University of Oslo, Oslo, Norway; National Research University Higher School of Economics, Moscow, Russia; Lomonosov Moscow State University, Moscow, Russia; Department of Tumor Biology, Institute for Cancer Research, Oslo University Hospital, Oslo, Norway; Center for Multimodal Imaging and Genetics, University of California San Diego, USA

## Abstract

Discoveries from genome-wide association studies often contain large clusters of highly correlated genetic variants, which makes them hard to interpret. In such cases, finemapping the underlying causal variants become important. Here we present a new method, the Finemap-MiXeR, based on a variational Bayesian approach for finemapping genomic data, i.e., determining the causal single nucleotide polymorphisms (SNPs) associated with a trait at a given locus after controlling for correlation among genetic variants due to linkage disequilibrium. Our approach is based on the optimization of Evidence Lower Bound of the likelihood function obtained from the MiXeR model. The optimization is done using Adaptive Moment Estimation Algorithm, allowing to obtain posterior probability of each SNP to be a causal variant. We tested Finemap-MiXeR in a range of different scenarios, using both synthetic and real data from the UK Biobank, using standing height phenotype as an example. In comparison to the existing finemapping methods FINEMAP and SuSiE methods, we observed that Finemap-MiXeR in most cases has better accuracy. Furthermore, it is computationally efficient, and unlike other methods the complexity is not increasing as the number of causal SNPs or the heritability increases. We show that our finemapping algorithm identifies a small number of genetic variants per locus which are informative for predicting the phenotype in an independent sample.

## Introduction

Genome-wide association studies (GWAS) have discovered hundreds of loci associated with complex human traits and disorders [1]. Such studies test for association between single nucleotide polymorphisms (SNPs) in a genotyped cohort, and the corresponding trait of interest (phenotype data). The results of a GWAS are available as summary statistics, which include z-scores and p-values for each SNP, and also may include the Linkage Disequilibrium (LD) matrix showing the correlations among SNPs. While traits are generally associated with many SNPs (with low p-values), most of these associated SNPs may not have a direct relation to the given trait [2]. In particular, a SNP may not be regarded as causal simply based on the GWAS summary statistics, since it may have a low p-value or a high z-score due to its high LD with another causal SNPs. Furthermore, causal SNPs may be missed in GWAS due to insufficient statistical power or unmeasured/unimputed SNPs [3]. Finemapping studies aim to identify causal SNPs, associated with a trait at a given locus after controlling for correlation among genetic variants due to LD, thus eliminating the SNPs that have indirect relation with the trait.

There are many existing finemapping methods in the literature. As stated in a review [4], Bayesian methods are useful to determine causal SNPs compared to other heuristic and penalized regression methods. Many Bayesian methods allow for more than one causal SNP per locus, and aim to infer causal SNP configuration under a given probabilistic model. Most of the early methods such as BIMBAM [5], CAVIAR [6], CAVIARBF [7], and PAINTOR [8] all rely on exhaustive searches of the possible causal configurations for a given locus and calculating corresponding posterior probabilities of being causal. Despite the accuracy of these methods, they are computationally inefficient, especially if the number of causal SNPs (k) or the total number of SNPs per locus (M) increases. Having multiple causal SNPs in a locus is a plausible situation that is often observed in different traits. For instance, it is shown that number of causal SNPs for prostate cancer in different regions ranges from 1 to 5 [9]. In particular, there are 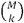 possible configurations that make exhaustive searches infeasible for large numbers of M and k. In order to tackle this problem, Benner et. al developed a computationally efficient method called FINEMAP [10] that performs calculation of the likelihood function using Cholesky Decomposition, and then search possible configurations via the Shotgun Stochastic Search [11]. Thanks to these improvements, the computational complexity has been reduced considerably, while preserving the same accuracy as the previous Bayesian exhaustive search methods like CAVIARBF. An extension of FINEMAP method which estimates the effect sizes and regional heritability under the probabilistic is also available [12].

Another recent approach to finemapping is based on applying a modified version of Single Effect Regression model, called the Sum of Single effects (SuSiE) [13]. The main idea behind this method is optimizing the SuSiE model by eliminating the effect of each causal SNP iteratively using Iterative Bayesian stepwise selection (IBSS). As a result, SuSiE provides the list of credible sets which include the list of possible causal SNPs with corresponding uncertainties that are used determine which variable to choose when SNPs are highly correlated. In this paper, it is also shown that optimizing SuSiE model with IBSS is equivalent to the optimization of a variational approximation to the posterior distribution under this model. Compared to the other existing Bayesian variable selection methods available in the literature such as [14] their proposed method is computationally less complex and more suitable for inference on highly correlated variables. Based on experiments conducted in [13], it was observed that SuSiE had better accuracy than the previously published methods. While the original SuSiE algorithm requires individual-level genome and phenotype data as input, it has been expanded to SuSiE Regression Summary Statistics (RSS) method using only summary statistics-level data [15]. SuSiE RSS yields a similar accuracy as the original SuSiE algorithm, at the same time reducing the computational complexity.

In the current paper, we present Finemap-MiXeR, a novel variational Bayesian method that aims to determine causal SNPs for a given locus. We are utilizing the MiXeR model [2], a causal mixture model which relies on a biologically plausible prior distribution of genetic effect sizes and can estimate the heritability, polygenicity and discoverability of a given trait, and the overlap between two phenotypes. Using MiXeR Model, we derive the likelihood of being causal using the summary statistics, and then optimize it using a Variational Bayesian approach particularly achieved by optimizing the Evidence Lower Bound (ELBO) of the likelihood function. We have calculated the derivatives of the ELBO function and optimize it using the Adaptive Moment Estimation (ADAM) Algorithm [16]. This method requires summary statistics and LD-matrix in order to obtain the posteriors of SNPs being causal.

Since FINEMAP and SuSiE have been demonstrated to have better performance compared to other existing methods in terms of accuracy and computational complexity, we compared our method with these two methods in different aspects. Based on experiments on synthetic data and real data, we observed that our method has an equivalent or better accuracy compared to FINEMAP and SuSiE, across a broad range of scenarios. Furthermore, unlike other methods, the computational complexity is linear with respect to the number of SNPs per locus (M), and is not increasing as number of causals (k) or heritability (*h*^*2*^) increases. While our method initially has O(M^2^) calculations per iteration, we have shown that it can be reduced to O(p_c_M) where p_c_ corresponds the number of principal components with highest eigenvalues of the LD matrix which covers most of the variation (see Supplementary Notes for details). Therefore, Finemap-MiXeR can also be considered computationally more efficient in the sense that in each iteration the required computation is only scalable with M.

## Results

For performance analysis, we have compared our Finemap-MiXeR method with FINEMAP and SuSiE. We have also compared MiXeR-PCA and SuSiE RSS in the “Computational Complexity” section, but omitted their performances on accuracy, since their performances in terms of accuracy were almost identical with MiXeR and SuSiE respectively. For comparison, we have used two different analyses: Simulation with synthetic data and application to real data.

### Simulation with Synthetic data

The first scheme is evaluation of the performances with synthetic genotype data created via the Hapgen2 tool [17], and simulating the phenotypes by arbitrarily choosing the actual causal SNPs and heritability. Given NxM genome matrix (G) for N “subjects” and M SNPs, a phenotype vector with a desired heritability (*h*^*2*^) can be obtained by randomly pre-assigning the causal SNPs with a vector *β* (where *β*_i_=1 if the SNP is causal and 0 otherwise) as

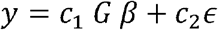

where 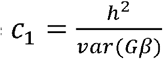 and 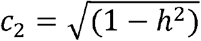 and ε represents non-genetic factors distributed by a standard normal distribution.

Using the procedure described above and sketched in Fig. 1, we have randomly chosen a locus from this synthetic genome data obtained the corresponding G matrix and determine the artificial phenotype vector y for different values of M and *h*^*2*^ for N=10 000. This procedure is repeated 50 times for different numbers of causals (k where 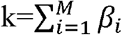). A basic quality control(QC) procedure has been applied to data, in line with previous studies, eliminating SNPs with MAF below 0.05, and randomly pruning one SNP from a pair with LD r2 correlation exceeding 0.99.

**Fig 1.**
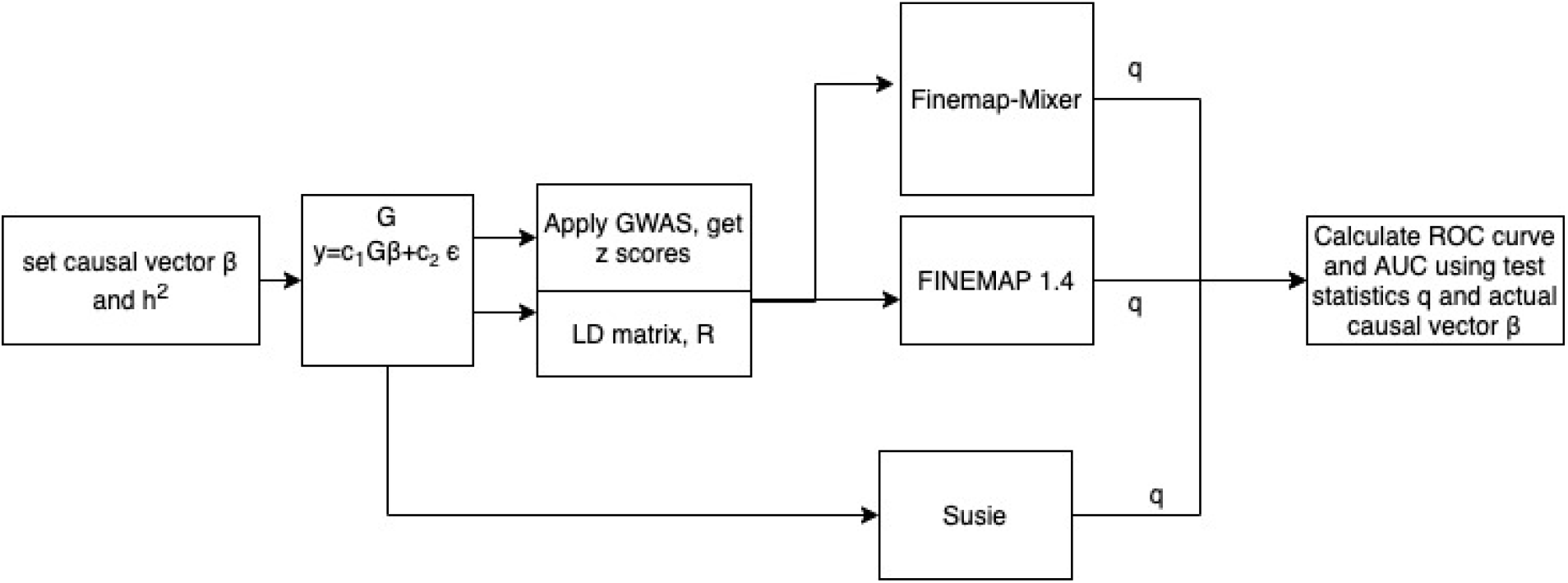
The Synthetic Data simulation procedure. Firstly, a random causal vector *β* and arbitrary heritability is determined. Then, a locus is randomly chosen from synthetic data (G) and desired phenotype (y) with given heritability is obtained using additive genetic model. Using G and y, z-scores are obtained by applying GWAS and then they are used as input for the tools to obtain posteriors of each SNP for being causal. Since we know ground truth causals (*β*), we then determine ROC curves for three methods and then calculate corresponding AUC.

Using posterior probabilities of being causal (q), we have evaluated the power to detect the actual causal SNPs and get corresponding Receiver Operating Characteristic curve for three methods, as shown in Fig. 2. Note that these values in the figures are the averaged values of 50 different experiments. As can be seen in Fig. 2, Finemap-MiXeR outperforms the other methods in different scenarios.

**Fig 2.**
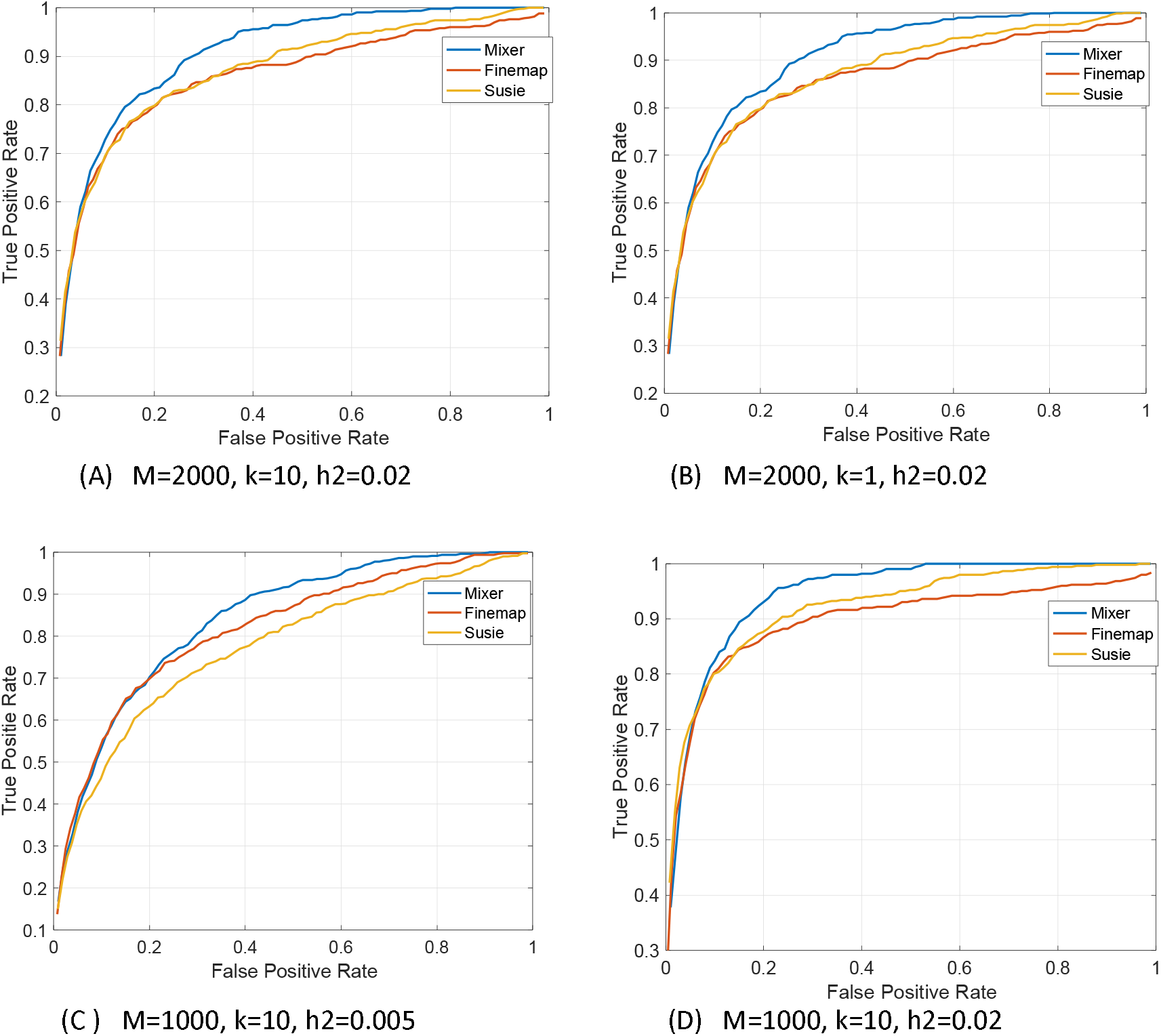
ROC curve comparison of three methods for different scenarios. The curves represent the average of 50 different simulations. M - number of SNPs per locus; k - number of causal variants; h2 - true SNP-based heritability per locus.

**Fig 3.**
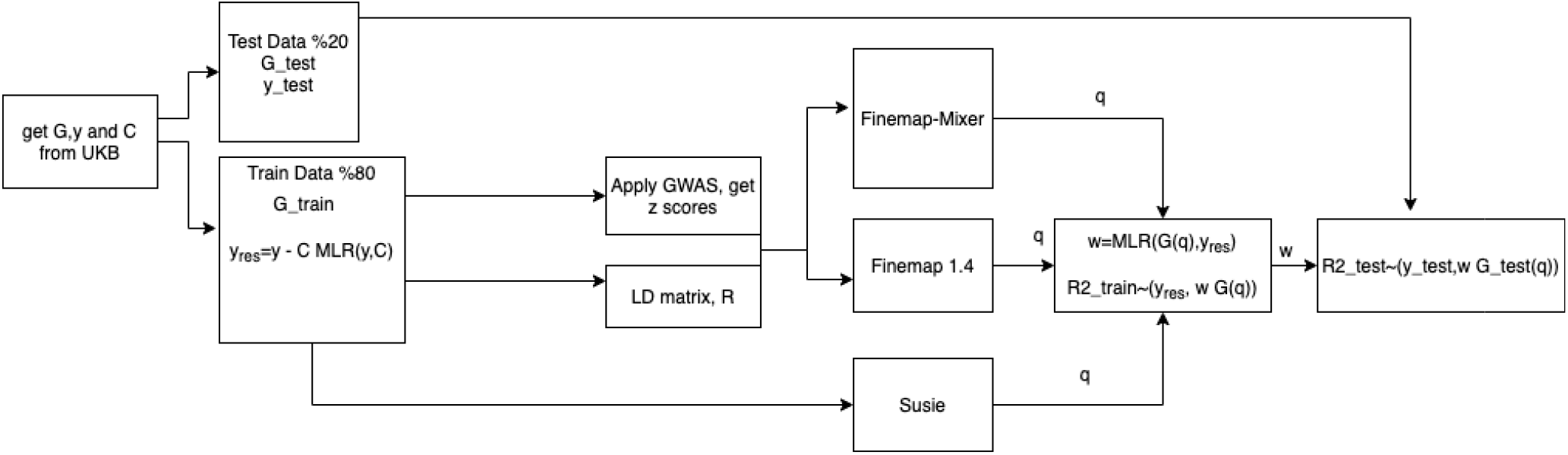
UKB data application procedure, using height as phenotype. We divided data into two as test and train. For train data, we firstly exclude the effects of covariates from phenotype and then follow the similar procedure to run tools as we did for synthetic data. Once we get posteriors, we then calculate obtain weight vector (w) of SNPs with highest posteriors using MLR and then use this vector to estimate the phenotype of test data.

The corresponding Area Under Curve (AUC) values for these ROC curves are given in Table 1. A broader overview of the performance comparison in terms of AUC can be examined in Table A in Supplementary Data. In this table, we have compared the performance of three methods using Area Under Curve (AUC) of ROC curves for 31 different scenarios. As can be observed from Table A, for most scenarios (21/31), Finemap-MiXeR outperforms the other methods in AUC. Although there are some cases where other methods are better (FINEMAP; 6/31, SuSiE 1/31), for those cases Finemap-MiXeR method has quite similar AUC values as compared to the best performing method and in three scenarios, the AUC values are same across methods. The mean AUC values of those 31 experiments for Finemap-MiXeR, SuSiE and FINEMAP are 0.932, 0.875 and 0.920, respectively. The corresponding median values are 0.961, 0.907 and 0.959. It can be worth to note that in some cases especially when the heritability is low, SuSiE may output all 0 vector as posterior for being causal.

**Table 1.** AUC values of three methods for different scenarios. Each scenario has been run via 50 times randomly. The best performing method is highlighted by star. An extended version of this table can be found at Table A in Supplementary Data.

| Nc | M | h2 | Tools | AUC |
| --- | --- | --- | --- | --- |
| 10 | 2000 | 0.020 | MiXeR* | 0.947 |
|  |  |  | Susie | 0.917 |
|  |  |  | FINEMAP | 0.901 |
| 1 | 2000 | 0.020 | MiXeR* | 0.989 |
|  |  |  | Susie | 0.981 |
|  |  |  | FINEMAP | 0.961 |
| 10 | 1000 | 0.005 | MiXeR* | 0.837 |
|  |  |  | Susie | 0.761 |
|  |  |  | FINEMAP | 0.806 |
| 10 | 1000 | 0.020 | MiXeR | 0.924 |
|  |  |  | Susie* | 0.907 |
|  |  |  | FINEMAP | 0.885 |

### Application to Real Data (UK Biobank Height)

We have used UK Biobank (UKB) genome data (N_total_=337 145 after QC) and standing height a phenotype to evaluate the performance of the Finemap-MiXeR method in real data. Since the ground truth causal variants for height are not known, we compared three methods by calculating polygenic prediction of height using the SNPs finemapped by these three algorithms, and then obtaining the correlation between the predicted height and the actual height. To achieve this, we split the individual-level UKB data into 80% for training, and 20% for testing. Once we obtain the SNPs and corresponding weights of those SNPs from train data to estimate the height, we then use these SNPs and their weights to estimate the height in the test data and evaluate the performance of three methods based on this estimation.

We conducted this procedure for multiple loci associated with height, varying in their heritability and loci length. In particular, we have chosen the loci that are strongly associated with height and then sorted them by their corresponding p-value of lead SNPs. We examined such 15 loci whose lead SNPs’ p-values are lowest. In order to get input data for the methods, we have applied GWAS to those loci and obtained z-scores. Then using these z-scores, we have run the 3 tools and got the posterior probabilities of being causal for each SNP.

Afterwards, we sorted these posteriors in descending order for each method and us corresponding SNPs to estimate height using Multiple Linear Regression (MLR). These SNP with highest posterior were eliminated as a stepwise manner. In particular, we excluded the SNPs which give higher p-value than a pre-defined threshold when adding them to MLR model to estimate height. Then we do the estimation of phenotypes using remaining (non-excluded) SNPs in MLR model.

For applying the procedure defined above to UKB data, firstly we have excluded covariates that are effective on height. Once we have extracted genotype (G) and phenotype data (y) from UKB, we have eliminated the effects of covariates (C) such as age, sex, and principal components. To achieve this, we first fit C and y using MLR and reduce the effect of C from phenotype as:

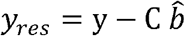

where y_res_ corresponds to a Covariates-free residual phenotype vector. In the simulations, we are using this vector and corresponding G matrix and apply GWAS to obtain z-scores, which is the input for FINEMAP and Finemap-MiXeR. For SuSiE, we are using *y*_*res*_ and G instead.

As stated above we have examined the loci whose lead SNPs have the lowest p-values. We run these three algorithms for these loci, assuming that there are 10 putative causal SNPs for each locus (note that we do not know how many causal SNPs we have in the real data) and get corresponding posterior (q) of being causal for each SNP. Afterwards, these posteriors are sorted in descending order we then apply a stepwise SNP selection procedure among those sorted SNPs by adding each of them to our SNP list, namely *q*_*list*_, if the existence of such a SNP leads to having a lower p-value than a pre-defined threshold (10^−2^) for the estimation of the phenotype with MLR. Then we get corresponding coefficients of the selected SNPs in *p*_*list*_, using G(*p*_*list*_) that is the columns of G indicated by q_list_ and *y*_*res*_ as

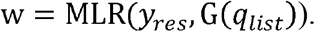

We have already split the actual data, using N=0.8 x N_total_ for training, and the rest for testing. In other words, w has been calculated using training data, and then it has been used to estimate the covariate-free test phenotype data y_test_ by w ×x G_test_(*q*_*list*_), followed by a comparison of the performance on estimation of the phenotypes by 3 methods. We calculated R2 of this estimation, and actual phenotype, as:

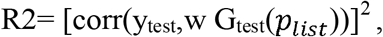

and used R2 metric to compare the performance of the methods.

As can be seen in Table 2, among the 15 scenarios that differ from chromosomes, loci length and heritability, in most cases (11/15), our method has the highest R2 value on test data, while SuSiE and FINEMAP performed better for three cases and one case, respectively. One of the important aspects of the results is that the number of SNPs used to predict y_test_ is generally higher when compared to other methods. In other words, our Finemap-MiXeR method is able to detect more causal SNPs than others in many cases and this leads to obtain better estimation for many cases. For instance, as given in Table 2, on chromosome 2, a locus whose lead SNP is rs78198962, Finemap-MiXeR method was able to detect 24 SNPs to predict phenotype, while other methods only detected 7 SNPs at this locus, and this yields that our method has a much higher R2 value than other methods.

**Table 2.** R2 value of phenotype (height) estimation for different locus in UKB data. For each locus (all of them are strongly associated with height and sorted by corresponding p value of lead SNPs), we have applied GWAS and obtained z-scores. Then using these z-scores, we have run the 3 tools and got the posterior probabilities of being causal for each SNP. Then, we used first 50 SNPs with highest posteriors for each method and use them for Multiple Linear Regression (MLR). These SNPs were eliminated as a stairwise manner. In particular, we excluded the SNPs which give higher p-value than a pre-defined threshold (10^−2^) when adding them to MLR and only use the rest to estimate the height. The p-value and R2 in this table correspond to p-value and R2 to model for train and test data for selected SNPs, while “# of SNPs selected” gives the total number of SNPs that is used in the ultimate MLR model. Heritability at given locus is given at h2 column and obtained using MLR. For each case, the best performing method in terms of R2 value for test data is highlighted by star.

| chr | lead SNP | p-value of the lead SNP | M | h2 | Tools | p-value train | R2 train | # SNPs selected | p-value test | R2 test |
| --- | --- | --- | --- | --- | --- | --- | --- | --- | --- | --- |
| 3 | rs2871960 | 1.25E-285 | 2878 | 0.01531 | Finemap-MiXeR | 1.737E-231 | 0.00391 | 11 | 1.58298E-51 | 0.0036383 |
|  |  |  |  |  | SuSiE* | 1.536E-231 | 0.00391 | 10 | 2.08141E-55 | 0.0036474 |
|  |  |  |  |  | FINEMAP | 8.52E-135 | 0.00227 | 5 | 7.01139E-37 | 0.0023893 |
| 11 | rs7952436 | 6.41E-153 | 6587 | 0.02924 | Finemap-MiXeR* | 2.221E-208 | 0.00352 | 23 | 3.5078E-60 | 0.00397 |
|  |  |  |  |  | SuSiE | 6.061E-176 | 0.00297 | 11 | 1.8302E-57 | 0.00379 |
|  |  |  |  |  | FINEMAP | 6.061E-176 | 0.00297 | 11 | 1.8302E-57 | 0.00379 |
| 18 | rs7235010 | 3.22E-144 | 3771 | 0.01940 | Finemap-MiXeR | 8.358E-241 | 0.00407 | 24 | 6.5221E-68 | 0.00450 |
|  |  |  |  |  | SuSiE* | 2.287E-237 | 0.00401 | 15 | 3.9051E-68 | 0.00451 |
|  |  |  |  |  | FINEMAP | 0.00023459 | 0.00005 | 1 | 0.06453989 | 0.00005 |
| 15 | rs72755233 | 1.86E-134 | 2737 | 0.01255 | Finemap-MiXeR* | 2.17E-121 | 0.00204 | 11 | 4.94547E-32 | 0.0023604 |
|  |  |  |  |  | SuSiE | 3.687E-133 | 0.00224 | 13 | 3.18455E-30 | 0.0019377 |
|  |  |  |  |  | FINEMAP | 9.0332E-53 | 0.00087 | 8 | 9.6229E-15 | 0.000891 |
| 6 | rs41271299 | 1.28E-139 | 542 | 0.00411 | Finemap-MiXeR* | 7.274E-121 | 0.00203 | 5 | 9.33652E-25 | 0.0015673 |
|  |  |  |  |  | SuSiE | 4.978E-123 | 0.00207 | 5 | 2.92425E-24 | 0.0015337 |
|  |  |  |  |  | FINEMAP | 1.2516E-07 | 0.0001 | 3 | 0.08364728 | 4.448E-05 |
| 19 | rs62621197 | 1.83E-134 | 213 | 0.00231 | Finemap-MiXeR* | 5.222E-91 | 0.00152 | 11 | 6.14402E-30 | 0.0020184 |
|  |  |  |  |  | SuSiE | 4.0278E-80 | 0.00133 | 7 | 3.93375E-30 | 0.0019315 |
|  |  |  |  |  | FINEMAP | 1.9097E-82 | 0.00137 | 8 | 3.6422E-30 | 0.0019338 |
| 7 | rs798528 | 2.90E-132 | 1224 | 0.00665 | Finemap-MiXeR* | 2.429E-118 | 0.00199 | 13 | 1.03864E-23 | 0.0014965 |
|  |  |  |  |  | SuSiE | 6.706E-120 | 0.00201 | 5 | 1.40762E-22 | 0.0014199 |
|  |  |  |  |  | FINEMAP | 7.1052E-14 | 0.00021 | 5 | 0.006312164 | 0.0001109 |
| 6 | rs9496369 | 3.26E-126 | 2020 | 0.00998 | Finemap-MiXeR | 1.955E-129 | 0.00217 | 13 | 1.14787E-27 | 0.0018644 |
|  |  |  |  |  | SuSiE* | 3.453E-139 | 0.00234 | 9 | 6.26433E-30 | 0.0019178 |
|  |  |  |  |  | FINEMAP | 8.8157E-36 | 0.00058 | 5 | 7.80823E-06 | 0.000297 |
| 17 | rs6505216 | 4.98E-122 | 3625 | 0.01714 | Finemap-MiXeR* | 8.578E-152 | 0.00256 | 21 | 2.13233E-32 | 0.0020852 |
|  |  |  |  |  | SuSiE | 0.00025073 | 5E-05 | 1 | 0.000209111 | 0.0002043 |
|  |  |  |  |  | FINEMAP | 4.111E-19 | 0.0003 | 6 | 2.93874E-07 | 0.0003907 |
| 13 | rs3118906 | 3.87E-113 | 2905 | 0.01492 | Finemap-MiXeR* | 1.803E-190 | 0.00301 | 20 | 1.03624E-37 | 0.0027457 |
|  |  |  |  |  | SuSiE | 1.813E-154 | 0.0026 | 9 | 3.8931E-38 | 0.0024746 |
|  |  |  |  |  | FINEMAP | 1.7445E-37 | 0.00261 | 5 | 4.34691E-12 | 0.0026126 |
| 20 | rs1884897 | 6.08E-101 | 1292 | 0.00739 | Finemap-MiXeR* | 9.391E-157 | 0.00264 | 18 | 6.28075E-27 | 0.0017144 |
|  |  |  |  |  | SuSiE | 7.451E-156 | 0.00262 | 13 | 2.4915E-26 | 0.0016739 |
|  |  |  |  |  | FINEMAP | 6.999E-153 | 0.00257 | 12 | 6.14739E-25 | 0.0015796 |
| 5 | rs244711 | 1.89E-98 | 1864 | 0.00992 | Finemap-MiXeR* | 6.234E-156 | 0.00263 | 19 | 3.36163E-29 | 0.0019384 |
|  |  |  |  |  | SuSiE | 6.887E-140 | 0.00235 | 11 | 7.33204E-29 | 0.0018454 |
|  |  |  |  |  | FINEMAP | 1.8253E-65 | 0.00108 | 9 | 5.33601E-13 | 0.0007737 |
| 2 | rs78198962 | 2.70E-58 | 3481 | 0.01666 | Finemap-MiXeR* | 1.721E-146 | 0.00307 | 24 | 1.77611E-39 | 0.00302 |
|  |  |  |  |  | SuSiE | 1.231E-121 | 0.00204 | 7 | 1.02703E-32 | 0.0021067 |
|  |  |  |  |  | FINEMAP | 1.231E-121 | 0.00204 | 7 | 1.02703E-32 | 0.0021067 |
| 5 | rs10805484 | 8.10E-48 | 2761 | 0.01191 | Finemap-MiXeR | 6.6539E-60 | 0.00099 | 11 | 1.91563E-08 | 0.0004692 |
|  |  |  |  |  | SuSiE | 2.831E-67 | 0.00112 | 10 | 2.59383E-08 | 0.0004605 |
|  |  |  |  |  | FINEMAP* | 3.7099E-62 | 0.00103 | 9 | 1.00019E-09 | 0.0005546 |
| 19 | rs12185519 | 1.02E-43 | 598 | 0.00327 | Finemap-MiXeR* | 1.1575E-44 | 0.00073 | 6 | 7.46048E-15 | 0.0008984 |
|  |  |  |  |  | SuSiE | 4.06E-49 | 0.00081 | 7 | 2.04492E-13 | 0.0008017 |
|  |  |  |  |  | FINEMAP | 1.5407E-14 | 0.00022 | 2 | 8.35238E-07 | 0.0003607 |

### Computational Complexity

The computational complexity of Finemap-MiXeR relies on the online calculation of the first derivatives of the ELBO of the likelihood ℒ (*q*, θ). As discussed in detail in Supplementary Notes, our Finemap-MiXeR method requires O(M^2^) computations per iteration. In Supplementary Notes, we have also shown that we can reduce the complexity from O(M^2^) to O(p_c_M) by preserving accuracy, where p_c_ << M. This improvement is achieved by using Principal Component Analysis (PCA) as described in the Supplementary Notes and p_c_ corresponds to first p_c_ principal components of the matrix of interest that is used to calculate the derivatives. We refer to this method as Finemap-MiXeR PCA. In SuSiE, the number of computations per iteration is O(kMN), and in its extension SuSiE RSS, it is O(kM^2^). In FINEMAP, the worst-case computation required per iteration is O(k^2^M). However, the algorithm was optimized to perform the search only among the SNPs with non-negligible posterior probabilities of being causal, using a hash table in order not to recalculate the same configurations. Thus, the complexity is expected to be reduced when the signal (heritability) is low. We have examined the computation time of Finemap-MiXeR, SuSiE and FINEMAP using the same data with different parameters. It is important to note that runtimes may largely differ due to different implementation (FINEMAP software used C++ code and is distributed as pre-compiled executable, SuSiE is an R package, Finemap-MiXeR is implemented using MATLAB). On the other hand, we can still compare how the runtime scales with respect to k, M, and h^2^ parameters. It is worth noting that the methods other than SuSiE are independent of N, since they use summary statistics, obviating the need to use N as a varying parameter. For comparison, synthetic data created by hapgen2 (N=10 000) are used.

As can be seen in Fig. 4, in Finemap-MiXeR, the required running time increases as the square of M. Similarly, for SuSiE RSS, it increases as the square of M, but it also scales linearly with k. In SuSiE, the running time is proportional to M and N and the running time is higher compared to SuSiE RSS when N<M, but when M increases, SuSiE is faster than SuSiE RSS as expected. On the other hand, the FINEMAP computation time increases directly proportional with M, but is more sensitive to the increase in h^2^ (which is an expected behavior as explained above). Furthermore, in SuSiE, SuSiE RSS and FINEMAP, the computational time increases as the number of causal variants increases, while in Finemap-MiXeR, the number of causal variants does not affect computational complexity. Finally, our extended version of Finemap-MiXeR, Finemap-MiXeR PCA, reduces the rate of increase of computation time as M increases. This is expected, since computation per iteration is proportional with p_c_M, where p_c_ is generally in the order of 100 at most. Although this method consumes some time to determine eigenvalues and eigenvectors before starting the iteration, it is still much faster than the Finemap-MiXeR and it reduces the rate of increase with M. As future work, we are planning to optimize these computations with more efficient methods to perform PCA.

**Fig 4.**
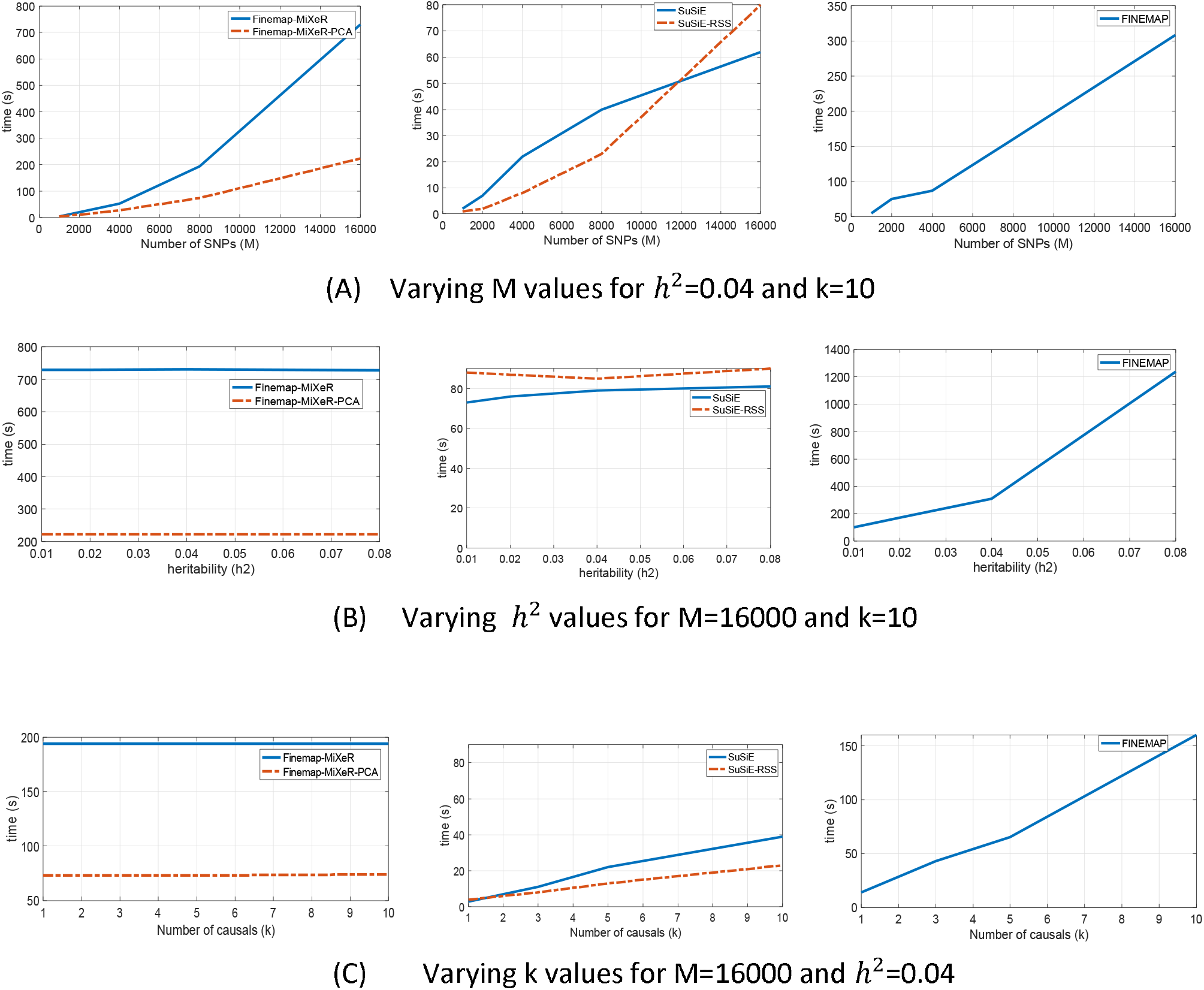
Run time comparison (in seconds) of the three methods, with their variants. Note that we have also included SuSiE RSS which uses SuSiE method with summary statistics, and Finemap-MiXeR PCA which reduces computational complexity by applying PCA. All tools are run in HPC with Intel Xeon CPU E5-2698 v4 @2.20GHZ. Unlike other methods, Finemap-MiXeR is only scalable with M and its computational complexity increases by M^2^. When we apply Finemap-MiXeR PCA, the computational complexity further reduces to the order of M.

## Discussion

We have developed a novel finemapping method, the Finemap-MiXeR, using the MiXeR model and a variational Bayesian approach. Our method relies on the optimization of the ELBO of the likelihood of the MiXeR model, using the ADAM algorithm. We have shown that the Finemap-MiXeR method has better accuracy and better computational performance compared to the existing methods.

The Finemap-MiXeR method performs better in terms of accuracy compared to other methods when we conduct comprehensive experiments on synthetic genomic data with different parameters (heritability, number of causals, loci length). The performance improvements were also observed in applications with real genomic data. To this end, for comparison, we have applied the methods on height, using samples from the UK Biobank. We evaluated multiple loci associated with height, varying in their heritability and loci length. The Finemap-MiXeR method outperformed the other methods in most scenarios, yielding better accuracy in estimating the phenotype. One of the main reasons of this performance improvements is MiXeR model’s flexibility in obtaining Bayesian inference for finemapping which leads to detect more causal variants than others in real applications.

Another benefit of Finemap-MiXeR is its computational effectiveness. Thanks to the MiXeR model with our tractable optimization function, our method’s complexity only depends on M, and does not increase as the number of causal and/or heritability increase, unlike the other methods do. In particular, although our method’s complexity is increased by O(M^2^) and thus is comparable with SuSiE (O(k M N)) and SuSiE RSS (O(k M^2^)), our method’s complexity is independent of the number of causals. Furthermore, unlike FINEMAP method, our method’s computational complexity is independent from the heritability. Finally, it is possible to reduce the complexity of our method to O(p_c_ M) by using Principal Component Analysis (PCA) to perform matrix vector multiplications.

Although we have assumed that all SNPs have equal priors in our model and we do the calculations accordingly, in our proposed setting, it is also possible to apply enriched priors to improve the performance. The MiXeR model provides novel opportunities for several applications in statistical genetics. We can also develop an optimization procedure for priors for performance improvement and this is left as a future work. In addition to the current finemapping application, it can also be used for gene set enrichment analysis. Combining these approaches could provide novel opportunities for improving functional analyses of GWAS findings. In conclusion, our results show that Finemap-MiXeR is a fast and accurate method for finemapping analysis of GWAS data from complex human traits.

## Data availability

The datasets analyzed during the current study are available for download from the following URLs: UK Biobank accessed via application 27412, https://bbams.ndph.ox.ac.uk/ams/ (upon application); Hapgen synthetic genome data, https://github.com/comorment/mixer/tree/main/reference/hapgen

## Code availability

Finemap-MiXeR software and a tutorial example on how to use it will be available online upon publication (https://github.com/comorment/FinemapMixer)

## Acknowledgements

The authors were funded by the Research Council of Norway (#324499, #324252, #273291, #223273, #276082), EU H2020 RealMent (#964874), EU H2020 CoMorMent (#847776) KG Jebsen Stiftelsen, South East Norway Health Authority (#2022-073). This research has been conducted using the UK Biobank Resource under Application Number 27412. This work also used the TSD (Tjeneste for Sensitive Data) facilities, owned by the University of Oslo, operated and developed by the TSD service group at the University of Oslo, IT-Department (USIT), using resources provided by UNINETT Sigma2 -the National Infrastructure for High Performance Computing and Data Storage in Norway. C.d.L. was funded by Hoffman-La Roche.

## Author’s contributions

B.C.A and O.F. conceived the study; B.C.A and O.F. pre-processed the data; B.C.A performed all analyses, with conceptual input from D.V., D.K., A.A.S., O.F., O.A.A. and A.M.D; B.C.A. and O.F. drafted the manuscript; all authors contributed to and approved the final manuscript.

## Online Methods

### Variational Bayesian Inference on Mixer Model

The Finemap-MiXeR method takes summary statistics and a scaled LD matrix as input and, using the MiXeR model, it outputs the probability of each SNP being causal (*q*_*i*_), alongside with the expectancy of effect size of the SNP (μ_*i*_) and uncertainty 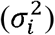 of the effect size estimate

In the MiXeR model, we can write the z-score of j-th SNP (*z*_*j*_) as a linear combination of the ground-truth effect sizes of all SNPs, and the coefficients comes from the scaled version of the LD matrix, namely A (see Supplementary Materials for the details of A). Given *a*_*ij*_ as the element of matrix A for i-th row and j-th column, we can write *z*_*j*_ as:

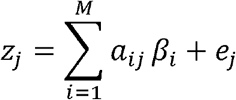

where *e*_*j*_ is error term and typically modelled as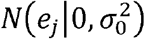

The MiXeR model assumes only a fraction of all SNPs (*π*_l_) in a locus are causal (i.e., have a non-zero ground-truth effect size *β* _*i*_) for a given phenotype, and can be postulated using a spike and slab prior as:

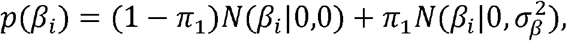

Where *π*_l_ ∈ [0,1]indicates the weight in the Gaussian mixture, *N* denotes a normal distribution (except for a special case *N* (.|0,0) which indicates probability mass at 0), and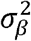 is a parameterof MiXeR model to represent the variance of non-zero effects and these parameters can be written as

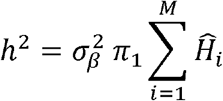

where 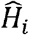is the estimated heterozygosity (See Supplementary materials for details).

For now, we assume that parameters 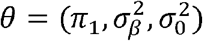 are the same across all SNPs, i.e.,do not depend on *i*. It is also possible to expand our proposed approach for SNPs with individual priors in future work on the model.

To determine the likelihood of the MiXeR model, we introduce the latent variables *u*_*i*_ ∈ {0,1} model We introduce the Lantent following Bernoulli distribution.*p*(*u*_*i*_) = *Bern* (*u*_*i*_ | *π*_1_). Then the full probabilistic Model is *p* (*z,β,u* | θ) = *p* (*z* |,*β*, θ) • *p* (β |*u*, θ … *p* (*u*| θ)) explicitly Written as;

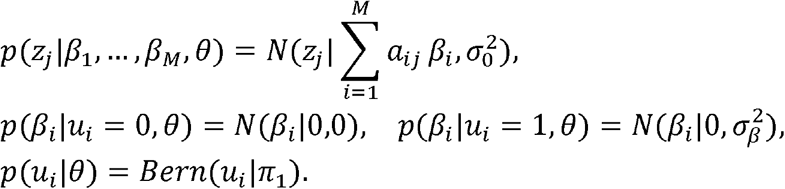

After observing *z* = (*z*_1_,…, *z*_*M*_)^T^, we can do inference on θ by maximum likelihood as;

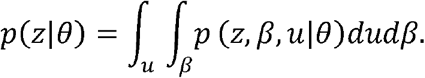

Note that *p*(*z |* θ) in its current form is neither tractable for optimization to achieve finemapping. In order to have a tractable optimization function, we use the Evidence Lower Bound (ELBO), instead of *p*(*z|* θ) to optimize its Variational Lower Bound as;

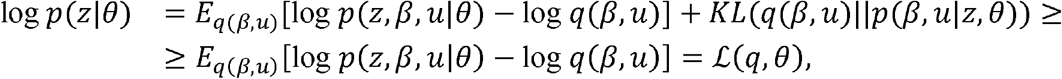

where ℒ (*q*, θ) is the variational lower bound of log *p*(*z|* θ) and *q*(*β,u*)is a distribution function from any parametric family. Choosing to be close to the *p*(*β,u | z*,θ)distribution leads to low values of the *KL* (*q* (*β,u*) | | *p*(*β,u | z*,θ) term, thus making ℒ (*q*, θ) a tight lower bound. In this situation the optimization of *p*(*z |* θwill be almost equivalent to the optimization of ℒ (*q*, θ) in a sense that any local maximum of the second problem will also yield local maximum of the original optimization problem). We will search *q*(*β,u*) from the following parametric family:

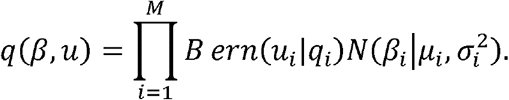

Using this model and parametric family, we can optimize ℒ (*q*, θ)and obtain the parameters of the *q* (*β, u*;) which corresponds to the posterior of the probability of being causal for each SNP (q_i_) and corresponding effect size μ *i*. In order to perform the optimization of ℒ (*q*, θ), we are using the Adaptive moment estimation (ADAM) algorithm, which computes the adaptive learning rate for each parameter using the first derivatives of ℒ (*q*, θ). For the details of the calculation of the derivatives and the ADAM algorithm, see Supplementary Notes.

### Hyperparameters

As stated before, we assumed that all hyperparameters of the MiXeR model 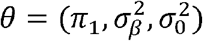 are the same across all SNPs. We left as a future work the optimization of those parameters using our variational Bayesian approach and, in this work, we use some fixed or pre-defined values instead. As suggested in [2], we chose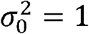, *π*_l_ is defined by user and 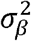 is estimated via using 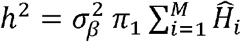 We had observed that choices of 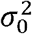, *π*_l_ and 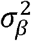do not lead tosignificant differences in the results but having individual *π*_l_ values for each SNP may lead a significant improvement. Although for the sake of simplicity, we omitted to optimize these parameters, we will focus on this problem in the future work.

## List of Supplementary Materials

1-Supplementary Notes: includes all the technical details of the proposed method.

2-Supplementary Table: includes more comprehensive results of the experiments related to synthetic data simulation

