## Supplementary Notes for "Finemap-MiXeR: A variational Bayesian approach for genetic finemapping"

#### Motivation

Human genetics studies show the individual genome of a person is linked to his/her observed features, or phenotypes, such as standing height, or the case/control status of heritable disorders such as schizophrenia. In this material, we consider quantitative phenotypes, $y \in \mathbb{R}$, but the results could be also extended to binary phenotypes $y \in \{0, 1\}$. The genome of a given person is typically encoded as a vector $\mathbf{g} = (g_1, \ldots, g_M)^T$, formed by $M$ genetic features, typically $g_i \in \{0, 1, 2\}$ indicates the number of reference alleles at a specific genetic locus, such as a single-nucleotide polymorphism (SNP).

Most of the available models assume a linear relationship between the genetic features and the observed phenotype, i.e. assume that there is a vector $\beta = (\beta_1, \ldots, \beta_M)^T$, such that

$$y = \mathbf{g}^T\beta + e = \sum_{i=1}^{M} \beta_i g_i + e,$$

where $\beta_i \in \mathbb{R}$ indicates the effect of $i$-th genetic variant, and $e$ is the residual contribution of non-genetic (environmental) factors, and non-linear genetic effects. Making inference about $\beta$ is crucial for our understanding of the genetics of complex human traits, as it may discover novel drug targets and suggest better treatment strategies.

Recent genome-wide association studies (GWAS) collect genotype and phenotype information from large cohorts of individuals. For example, UK Biobank study has released information for $N = 500.000$ individuals, $M = 93.000.000$ genetic variants, and tens of thousands of quantitative and binary phenotypes. To study a phenotype $y$, e.g. standing height, we have measurements $\mathbf{y} = (y_1, \ldots, y_N)^T$ across all individuals, and a genotype matrix $G = \{g_{ki}\}$, $k = 1, \ldots N$, $i = 1, \ldots M$. Our goal is to solve a system of linear equations $\mathbf{y} = G\beta$, with known $\mathbf{y}$ and $G$, w.r.t. unknown $\beta$.

As both $\mathbf{y}$ and $G$ contain sensitive (individual-level) information, GWAS studies typically keep this information confidential and release the *GWAS summary statistics*. Ignoring all genetic variants except $j$-th, consider a simple linear regression model $y \sim g_j$. The corresponding regression coefficient $\hat{\beta}_j$ can be estimated as $\hat{\beta}_j = \frac{\mathbf{v_j^T y}}{\mathbf{v_j^T v_j}}$, where $\mathbf{v}_j = (g_{1j}, \ldots, g_{Nj})^T$ is a vector of $j$-th genetic features across individuals. The standard error of $\hat{\beta}_j$ estimate vary substantially across genetic features, therefore in addition to $\hat{\beta}_j$ GWAS summary statistics provide statistical significance, $z_j = \frac{\hat{\beta}_j}{se(\beta_j)}$. It would be more correct to use the notation $t_j$ instead of $z_j$ and call it Wald's t-statistic, but due to large $N$ we usually denote it $z_j$ and refer to it as z-score. Across genetic variants $j = 1, \ldots, M$, we have a vector $\mathbf{z} = (z_1, \ldots, z_M)$ of z-scores, $z_j = \frac{\hat{\beta}_j}{se(\beta_j)}$. It is possible to derive that such $z$-scores are linearly related to $\beta$ via matrix $A = (a_{ij})$, derived from $G = (g_{ij})$:

$$z_j = \sum_{i=1}^{M} a_{ij}\beta_i + e_j,$$

where $p(e_j) = N(e_j|0, \sigma_0^2)$ and typically $\sigma_0^2 = 1$. The elements of matrix $A$ can be obtained by $a_{ij} = \sum_{i=1}^{M} \sqrt{N\hat{H}_i}\hat{r}_{ji}$ as defined in [1] where $\hat{H}_i$ is the sample heterozygosity of variant $i$ and $\hat{r}_{ji}$ is sample correlation coefficients between variant $i$ and $j$. The main goal is to solve a system of linear equations $\mathbf{z} = A\beta$, with known $\mathbf{z}$ and $A$, w.r.t. unknown $\beta$, knowing that $A$ is sparse and band matrix.

##### MiXeR Model

In MiXeR model, we postulate a spike-and-slab prior distribution on $\beta_i$:

$$p(\beta_i) = (1 - \pi_1)N(\beta_i|0, 0) + \pi_1 N(\beta_i|0, \sigma_\beta^2),$$

where $\pi_1 \in [0, 1]$ indicates the weight in the mixture, $N(\beta_i|0, \sigma_\beta^2)$ denotes the normal distribution function of $\beta_i$ with zero mean and $\sigma_\beta^2$ variance (except for a special case $N(\beta_i|0, 0)$ which indicates probability mass at 0), and $\sigma_\beta^2$ corresponds to the variance of non-zero effects and can be obtained from heritability($h^2$) as $h^2 = \sigma_\beta^2 \pi_1 \sum_{i=1}^M \hat{H}_i$. For now we assume that parameters $\theta = (\pi_1, \sigma_\beta^2, \sigma_0^2)$ are the same across all SNPs, i.e. do not depend on $i$. It is also possible to do the same analysis with SNP-specific priors but left as a future work.

We introduce latent variables $u_i \in \{0, 1\}$ following Bernoulli distribution, $p(u_i) = Bern(u_i|\pi_1)$ where $u_i = 1$ implies that SNP $i$ is causal and $u_i = 0$ otherwise. Then the full probabilistic model is $p(z, \beta, u|\theta) = p(z|\beta, \theta) \cdot p(\beta|u, \theta) \cdot p(u|\theta)$ and its product components can be written as:

$$\begin{aligned}
&p(z_j|\beta_1, \dots, \beta_M, \theta) = N\Big(z_j\Big|\sum_{i=1}^M a_{ij}\beta_i, \sigma_0^2\Big),\\
&p(\beta_i|u_i = 0, \theta) = N(\beta_i|0, 0), \quad p(\beta_i|u_i = 1, \theta) = N(\beta_i|0, \sigma_\beta^2),\\
&p(u_i|\theta) = Bern(u_i|\pi_1).
\end{aligned}$$

##### Variational Bayesian inference

A tricky part of the model is that $z_j$ may depend on multiple $\beta_i$. After observing $z = (z_1, \dots, z_M)^T$, we are aiming to do inference on $\theta$ by max. likelihood:

$$p(z|\theta) = \int_u \int_\beta p(z, \beta, u|\theta) du d\beta \to \max_\theta .$$

Next, for obtaining a tractable optimization function, we use Evidence Lower Bound (ELBO) and, instead of $p(z|\theta) \to \max_\theta$ and optimize its Variational Lower Bound:

$$\begin{aligned}
\log p(z|\theta) = E_{q(\beta,u)}[\log p(z, \beta, u|\theta) - \log q(\beta, u)] + KL(q(\beta, u)||p(\beta, u|z, \theta)) \geq\\
\geq E_{q(\beta,u)}[\log p(z, \beta, u|\theta) - \log q(\beta, u)] = \mathcal{L}(q, \theta) \to \max_{q,\theta},
\end{aligned}$$

where $KL(q(\beta, u)||p(\beta, u|z, \theta))$ is Kullback–Leibler divergence and it is a measure of how distribution $q(\beta, u)$ is different from $p(\beta, u|z, \theta)$. Therefore, choosing $q(\beta, u)$ close to the distribution of $p(\beta, u|z, \theta)$ leads to low values of $KL(q(\beta, u)||p(\beta, u|z, \theta))$ term, thus making $\mathcal{L}(q, \theta)$ a tight bound of $\log p(z|\theta)$. In this case, the optimization problems $p(z|\theta) \to \max_\theta$ and $\mathcal{L}(q, \theta) \to \max_{q,\theta}$ are almost equivalent (in a sense that any local maximum of the second problem will also yield a local maximum of the original optimization problem). We will search $q(\beta, u)$ from the following a parametric family:

$$q(\beta, u) = \prod_{i=1}^{M} Bern(u_i|q_i) N(\beta_i|\mu_i, \sigma_i^2).$$

To optimize $\mathcal{L}(q, \theta)$, we use ADAM algorithm [2] to explicitly optimize $q_i$, $\mu_i$ and $\sigma_i^2$ parameters as well as $\theta$. As you will observe in the next chapters, in order to apply ADAM algorithm, the first derivatives of the objective function are required. To do so, we need to obtain a tractable formula for $\mathcal{L}(q, \theta)$ which can be written initially as;

$$\begin{aligned}
\mathcal{L}(q, \theta) &= E_{q(\beta,u)} \log p(z|\beta, \theta) + E_{q(\beta,u)} \log \frac{p(\beta|u, \theta)p(u|\theta)}{q(\beta, u)} = \\
&= E_{q(\beta)} \log p(z|\beta, \theta) + E_{q(\beta)q(u)} \log \frac{p(\beta|u, \theta)p(u|\theta)}{q(\beta)q(u)} = \\
&= E_{q(\beta)} \log p(z|\beta, \theta) + E_{q(\beta)q(u)} \log \frac{p(\beta|u, \theta)}{q(\beta)} + E_{q(\beta)q(u)} \log \frac{p(u|\theta)}{q(u)} = \\
&= E_{q(\beta)} \log p(z|\beta, \theta) + E_{q(u)q(\beta)} \log \frac{p(\beta|u, \theta)}{q(\beta)} + E_{q(u)} \log \frac{p(u|\theta)}{q(u)} = \\
&= E_{q(\beta)} \log p(z|\beta, \theta) - E_{q(u)} KL(q(\beta)||p(\beta|u, \theta)) - KL(q(u)||p(u|\theta)) = \\
&= E_{q(\beta)} \log p(z|\beta, \theta) - E_{q(u)} \sum_{i=1}^{M} KL(q(\beta_i)||p(\beta_i|u_i, \theta)) - \sum_{i=1}^{M} KL(q(u_i)||p(u_i|\theta)).
\end{aligned}$$

**Calculating the derivatives of Variational Lower Bound**

In this section, we will obtain the first derivatives with respect to decision variables which are $\mu_i$, $\sigma_i^2$, and $q_i$ to optimize $\mathcal{L}(q, \theta)$. Firstly, we need to expand the terms of $\mathcal{L}(q, \theta)$ in order to ease derivative calculation. Assuming $z_j$'s are independent, $E_{q(\beta)} \log p(z|\beta, \theta)$ can be rewritten as $E_{q(\beta)} \log p(z|\beta, \theta) = E_{q(\beta)} \sum_{j=1}^{M} \log p(z_j|\beta, \theta)$ hence $\mathcal{L}(q, \theta)$ can also be represented as

$$\begin{aligned}
\mathcal{L}(q, \theta) = &\underbrace{E_{q(\beta)} \sum_{j=1}^{M} \log p(z_j|\beta, \theta)}_{T_1} - \underbrace{E_{q(u)} \sum_{i=1}^{M} KL(q(\beta_i)||p(\beta_i|u_i, \theta))}_{T_2} - \underbrace{\sum_{i=1}^{M} KL(q(u_i)||p(u_i|\theta))}_{T_3} = \\
&T_1 - T_2 - T_3.
\end{aligned} \tag{1}$$

Hence we divide $\mathcal{L}(q, \theta)$ into three terms in order to ease the workload for derivative calculation.

To deal with the derivatives of $E_{q(\beta)} \log p(z|\beta,\theta)$ (hence $T_1$) we need to employ reparametrization trick [3]. In particular, it is easy to compute $\frac{\partial E_{q(a)} f(b)}{\partial b}$, but it's unclear how to compute $\frac{\partial E_{q(a)} f(b)}{\partial a}$ and we have such cases in $T_1$. The Reparametrization trick allows us to circumvent this issue by using a parametric standard distribution $\epsilon$ (in our case, the standard normal distribution can be used for this purpose), and reformulating the function with this parametric function as

$$E_{q(\beta_i|\mu_i,\sigma_i^2)} \log p(z|\beta_i,\theta) = E_\epsilon \log p(z|\beta_i(\epsilon,\mu_i,\sigma_i^2),\theta).$$

Let $\boldsymbol{\epsilon} \in [\epsilon_1 \epsilon_2 \dots \epsilon_M] \sim \boldsymbol{N(0,I)}$, then reparametrization trick can be applied as

$$E_{q(\beta)} \sum_{j=1}^{M} \log p(z_j|\beta,\theta) =$$

$$E_{\boldsymbol{\epsilon}} \sum_{j=1}^{M} \log p(z_j|\beta_1 = \mu_1 + \sigma_1\epsilon_1, \beta_2, \dots \beta_i = \mu_i + \sigma_i\epsilon_i \dots \beta_M, \theta) \equiv E_{\boldsymbol{\epsilon}} \sum_{j=1}^{M} \log p(z_j|\beta = \mu + \sigma\boldsymbol{\epsilon}, \theta). \tag{2}$$

Since $z_j = \sum_{i=1}^{M} a_{ij}\beta_i + e_j$, then we may write its distribution as

$$z_{j|\beta=\mu+\sigma\boldsymbol{\epsilon},\theta} = \sum_{i=1}^{M} a_{ij}(\mu_i + \sigma_i\epsilon_i) + e_j \sim N(z_j|\sum_{i=1}^{M} a_{ij}(\mu_i + \sigma_i\epsilon_i), \sigma_0^2). \tag{3}$$

In order to calculate the gradients of $T_1$ (hence $E_{\boldsymbol{\epsilon}} \sum_{j=1}^{M} \log p(z_j|\beta = \mu + \sigma\boldsymbol{\epsilon}, \theta)$), it would be beneficial to write down $\log p(z_j|\beta = \mu + \sigma\boldsymbol{\epsilon}, \theta)$ explicitly. As Eq. 3 implies, it can be written as

$$\log p(z_j|\beta = \mu + \sigma\boldsymbol{\epsilon}, \theta) = -\log\left[\sqrt{2\pi\sigma_0^2}\right] - \frac{\left(z_j - \sum_{i=1}^{M} a_{ij}(\mu_i + \sigma_i\epsilon_i)\right)^2}{2\sigma_0^2}. \tag{4}$$

If we plug this expression into $T_1$, then it is possible to get the following expression:

$$\mathrm{T}_1 = E_{\boldsymbol{\epsilon}} \sum_{j=1}^{M} \log p(z_j|\beta = \mu + \sigma\boldsymbol{\epsilon}, \theta) = E_{\boldsymbol{\epsilon}} \sum_{j=1}^{M} -\log\left[\sqrt{2\pi\sigma_0^2}\right] - \frac{\left(z_j - \sum_{i=1}^{M} a_{ij}(\mu_i+\sigma_i\epsilon_i)\right)^2}{2\sigma_0^2}$$

$$= - \sum_{j=1}^{M} E_{\boldsymbol{\epsilon}} \log\left[\sqrt{2\pi\sigma_0^2}\right] - \sum_{j=1}^{M} E_{\boldsymbol{\epsilon}} \frac{\left(z_j - \sum_{i=1}^{M} a_{ij}(\mu_i+\sigma_i\epsilon_i)\right)^2}{2\sigma_0^2}$$

$$= -M\log\left[\sqrt{2\pi\sigma_0^2}\right] - \sum_{j=1}^{M} E_{\boldsymbol{\epsilon}} \frac{\left(z_j - \sum_{i=1}^{M} a_{ij}\mu_i - \sum_{i=1}^{M} a_{ij}\sigma_i\epsilon_i\right)^2}{2\sigma_0^2}$$

$$= -M\log\left[\sqrt{2\pi\sigma_0^2}\right] - \sum_{j=1}^{M} E_{\boldsymbol{\epsilon}} \frac{(z_j - \sum_{i=1}^{M} a_{ij}\mu_i)^2 - 2(z_j - \sum_{i=1}^{M} a_{ij}\mu_i)(\sum_{i=1}^{M} a_{ij}\sigma_i\epsilon_i) + (\sum_{i=1}^{M} a_{ij}\sigma_i\epsilon_i)^2}{2\sigma_0^2}.$$

If we expand the $E_\epsilon$ operator in $T_1$, we can deduce that the middle term vanishes since it has $E_\epsilon[\epsilon_i]$ as a product term;

$$E_\epsilon[-2(z_j - \sum_{i=1}^{M} a_{ij}\mu_i)(\sum_{i=1}^{M} a_{ij}\sigma_i\epsilon_i)] = -2(z_j - \sum_{i=1}^{M} a_{ij}\mu_i)(\sum_{i=1}^{M} a_{ij}\sigma_i E_\epsilon[\epsilon_i]) = 0, \tag{5}$$

and similarly, we can calculate the first term of $T_1$ as

$$E_\epsilon[(z_j - \sum_{i=1}^{M} a_{ij}\mu_i)^2] = (z_j - \sum_{i=1}^{M} a_{ij}\mu_i)^2. \tag{6}$$

The third term of $T_1$ is the most challenging part. Firstly we need to rewrite this term as the square of the summation

$$\left(\sum_{i=1}^{M} a_{ij}\sigma_i\epsilon_i\right)^2 = \sum_{i=1}^{M}\left((a_{ij}\sigma_i\epsilon_i)^2 + 2\sum_{k=i+1}^{M}(a_{kj}\sigma_k\epsilon_k a_{ij}\sigma_i\epsilon_i)\right) \tag{7}$$

and then if we employ $E_\epsilon$ operator we may get;

$$E_\epsilon\left(\sum_{i=1}^{M} a_{ij}\sigma_i\epsilon_i\right)^2 = \sum_{i=1}^{M}\left(E_\epsilon(a_{ij}\sigma_i\epsilon_i)^2 + 2E_\epsilon\sum_{k=i+1}^{M}(a_{kj}\sigma_k a_{ij}\sigma_i\epsilon_k\epsilon_i)\right), \tag{8}$$

which implies

$$E_\epsilon\left(\sum_{i=1}^{M} a_{ij}\sigma_i\epsilon_i\right)^2 = \sum_{i=1}^{M}\left((a_{ij}^2\sigma_i^2 E_\epsilon[\epsilon_i^2]) + 2\sum_{k=i+1}^{M}(a_{kj}\sigma_k a_{ij} E_\epsilon[\sigma_i\epsilon_k\epsilon_i])\right). \tag{9}$$

Note that $E_\epsilon[\epsilon_i^2] = 1$ and $E_\epsilon[\sigma_i\epsilon_k\epsilon_i)] = 0$ since $k > i$. Then it is possible to further simplify it as;

$$E_\epsilon\left(\sum_{i=1}^{M} a_{ij}\sigma_i\epsilon_i\right)^2 = \sum_{i=1}^{M}\left(a_{ij}^2\sigma_i^2\right). \tag{10}$$

Then $T_1$ can be written as;

$$
\begin{aligned}
T_1 =& - M \log\left[\sqrt{2\pi\sigma_0^2}\right] - \frac{1}{2\sigma_0^2}\sum_{j=1}^{M}\sum_{i=1}^{M}\left(a_{ij}^2\sigma_i^2\right) - \frac{1}{2\sigma_0^2}\sum_{j=1}^{M}(z_j - \sum_{i=1}^{M} a_{ij}\mu_i)^2 \\
=& -M \log\left[\sqrt{2\pi\sigma_0^2}\right] - \frac{T_A}{2\sigma_0^2} \qquad (11)
\end{aligned}
$$

where $T_A = \sum_{j=1}^{M}\sum_{i=1}^{M}\left(a_{ij}^2\sigma_i^2\right) + \sum_{j=1}^{M}(z_j - \sum_{i=1}^{M} a_{ij}\mu_i)^2$.

Note that, since our main motivation is obtaining $\frac{\partial \mathcal{L}_{q,\theta}}{\partial \mu_i}$, $\frac{\partial \mathcal{L}_{q,\theta}}{\partial \sigma_i^2}$, $\frac{\partial \mathcal{L}_{q,\theta}}{\partial q_i}$, $\frac{\partial \mathcal{L}_{q,\theta}}{\partial \theta}$. It is always worthwhile to keep in mind that $E_{\boldsymbol{\epsilon}}[\epsilon_i] = 0$ and $E_{\boldsymbol{\epsilon}}[\epsilon_i^2] = 1$. Then we may start by evaluating $\frac{\partial T_1}{\partial \mu_i}$:

$$
\begin{aligned}
\frac{\partial T_1}{\partial \mu_i} =& - \frac{\partial}{\partial \mu_i} M \log\left[\sqrt{2\pi\sigma_0^2}\right] \\
& - \frac{\partial}{\partial \mu_i}\sum_{j=1}^{M} E_{\boldsymbol{\epsilon}} \frac{(z_j - \sum_{i=1}^{M} a_{ij}\mu_i)^2 - 2(z_j - \sum_{i=1}^{M} a_{ij}\mu_i)(\sum_{i=1}^{M} a_{ij}\sigma_i\epsilon_i) + (\sum_{i=1}^{M} a_{ij}\sigma_i\epsilon_i)^2}{2\sigma_0^2} \\
=& - \sum_{j=1}^{M} E_{\boldsymbol{\epsilon}} \frac{\partial}{\partial \mu_i} \frac{(z_j - \sum_{i=1}^{M} a_{ij}\mu_i)^2 - 2(z_j - \sum_{i=1}^{M} a_{ij}\mu_i)(\sum_{i=1}^{M} a_{ij}\sigma_i\epsilon_i) + (\sum_{i=1}^{M} a_{ij}\sigma_i\epsilon_i)^2}{2\sigma_0^2}.
\end{aligned}
\qquad (12)
$$

In Eq. 12, the rightmost term vanishes since it is independent of $\mu_i$ and the middle term vanishes since it includes $E_{\boldsymbol{\epsilon}}[\epsilon_i] = 0$. Therefore, it is possible to simplify the expression as

$$
\frac{\partial T_1}{\partial \mu_{i^*}} = - \sum_{j=1}^{M} E_{\boldsymbol{\epsilon}} \frac{\partial}{\partial \mu_i} \frac{(z_j - \sum_{i=1}^{M} a_{ij}\mu_i)^2}{2\sigma_0^2} = - \sum_{j=1}^{M} \frac{\partial}{\partial \mu_i} \frac{(z_j - \sum_{i=1}^{M} a_{ij}\mu_i)^2}{2\sigma_0^2} \qquad (13)
$$

$$
= \frac{1}{\sigma_0^2}\sum_{j=1}^{M} a_{i^*j}(z_j - \sum_{i=1}^{M} a_{ij}\mu_i). \qquad (14)
$$

In a similar manner, it is possible to calculate $\frac{\partial T_1}{\partial \sigma_i}$:

$$
\frac{\partial T_1}{\partial \sigma_i} = - \sum_{j=1}^{M} E_{\boldsymbol{\epsilon}} \frac{\partial}{\partial \sigma_i} \frac{(z_j - \sum_{i=1}^{M} a_{ij}\mu_i)^2 - 2(z_j - \sum_{i=1}^{M} a_{ij}\mu_i)(\sum_{i=1}^{M} a_{ij}\sigma_i\epsilon_i) + (\sum_{i=1}^{M} a_{ij}\sigma_i\epsilon_i)^2}{2\sigma_0^2}. \qquad (15)
$$

Similar to the previous derivation, one can observe that the first and the second term are vanished by the derivative and expectation operators respectively. Hence the expression can be further simplified as

$$\frac{\partial T_1}{\partial \sigma_{i^*}} = -\sum_{j=1}^{M} \frac{\partial}{\partial \sigma_i^*} E_{\boldsymbol{\epsilon}} \frac{(\sum_{i=1}^{M} a_{ij}\sigma_i\epsilon_i)^2}{2\sigma_0^2} = -\sum_{j=1}^{M} E_{\boldsymbol{\epsilon}} \frac{\partial}{\partial \sigma_i^*} \frac{(\sum_{i=1}^{M} a_{ij}\sigma_i\epsilon_i)^2}{2\sigma_0^2} \tag{16}$$

$$= -\sum_{j=1}^{M} E_{\boldsymbol{\epsilon}} 2a_{i^*j}\epsilon_i^* \frac{(\sum_{i=1}^{M} a_{ij}\sigma_i\epsilon_i)}{2\sigma_0^2}. \tag{17}$$

Note that the purpose of proposing $i^*$ is to avoid confusion with the summation index in Eq. 16. Once we have eliminated this summation, then we will replace it as $i$. Since $\epsilon_i$s are independent and identically distributed, $E_{\boldsymbol{\epsilon}}[\epsilon_i^*\epsilon_i] = 1$ iff $i = i^*$ and 0 otherwise. Using these, we may obtain the following expression

$$\frac{\partial T_1}{\partial \sigma_{i^*}} = -\frac{1}{2\sigma_0^2} \sum_{j=1}^{M} 2a_{i^*j}^2 \sigma_{i^*}. \tag{18}$$

Then using the chain rule $\frac{\partial T_1}{\partial \sigma_{i^*}} = \frac{\partial T_1}{\partial \sigma_{i^*}^2} \frac{\partial \sigma_{i^*}^2}{\partial \sigma_{i^*}}$, we can easily obtain $\frac{\partial T_1}{\partial \sigma_{i^*}^2}$ as

$$\frac{\partial T_1}{\partial \sigma_{i^*}^2} = \frac{-1}{2\sigma_{i^*}} \frac{1}{2\sigma_0^2} \sum_{j=1}^{M} 2a_{i^*j}^2 \sigma_{i^*} = \frac{-1}{4\sigma_0^2} \sum_{j=1}^{M} 2a_{i^*j}^2. \tag{19}$$

It is also quite straightforward to calculate the gradient of $T_1$ with respect to $\theta$ which is $\nabla_\theta T_1 = \begin{bmatrix} \dfrac{\partial T_1}{\partial \pi_1} \\ \dfrac{\partial T_1}{\partial \sigma_\beta^2} \\ \dfrac{\partial T_1}{\partial \sigma_0^2} \end{bmatrix}$.

Since $T_1$ only depends on $\sigma_0^2$ among variables of $\theta$, we may readily calculate it by utilizing (11) as

$$\frac{\partial T_1}{\partial \sigma_0^2} = \frac{T_A - M\sigma_0^2}{2\sigma_0^4}. \tag{20}$$

Hence,

$$\nabla_\theta T_1 = \begin{bmatrix} 0 \\ 0 \\ \frac{T_A - M\sigma_0^2}{2\sigma_0^4} \end{bmatrix}. \tag{21}$$

For $T_2$, note that it is possible to rewrite it as

$$E_{q(u)} \sum_{i=1}^{M} KL(q(\beta_i)||p(\beta_i|u_i, \theta)) = \sum_{i=1}^{M} E_{q(u)} KL(q(\beta_i)||p(\beta_i|u_i, \theta)). \tag{22}$$

Then, for $u_i = 0$ and $u_i = 1$ we write the expectation explicitly as:

$$\begin{aligned} E_{q(u)} KL(q(\beta_i)||p(\beta_i|u_i, \theta)) &= q(u_i = 0) KL(q(\beta_i)||N(0, \delta^2)) \\ &+ q(u_i = 1) KL(q(\beta_i)||N(0, \sigma_\beta^2)) \\ &= (1 - q_i) KL(q(\beta_i)||N(0, \delta^2)) + q_i KL(q(\beta_i)||N(0, \sigma_\beta^2)) \end{aligned}$$

where $\delta^2$ is a sufficiently small and adjustable parameter to approximate $N(0, \delta^2)$ as Dirac delta function.

In order to obtain the gradients of $T_2$, the first step would be representing $q(\beta_i)$ and $q(u_i)$ explicitly. Using $q(\beta, u)$, it is possible to write them as

$$\begin{aligned} q(\beta_i) &= N(\beta_i|\mu_i, \sigma_i^2) \\ q(u_i) &= Bern(u_i|q_i). \end{aligned}$$

Then it is possible to evaluate KL divergence in $T_2$ (and also $T_3$). In particular, $T_2$ involves KL divergence of two normal distributions and can be written as [1]

$$\begin{aligned} T_2 = \sum_{i=1}^{M} E_{q(u)} KL(q(\beta_i)||p(\beta_i|u_i, \theta)) = \sum_{i=1}^{M} (1 - q_i) \left( \log(\frac{\delta}{\sigma_i}) + \frac{\sigma_i^2 + \mu_i^2}{2\delta^2} - \frac{1}{2} \right) + \\ \sum_{i=1}^{M} q_i \left( \log(\frac{\sigma_\beta}{\sigma_i}) + \frac{\sigma_i^2 + \mu_i^2}{2\sigma_\beta^2} - \frac{1}{2} \right). \end{aligned}$$

Then it is straightforward to calculate derivatives as:

[1] Let, x and y be two normal distributions with means $\mu_1$, $\mu_2$ and standard deviations $\sigma_1$, $\sigma_2$, respectively. Then $KL(x||y) = \left( \log(\frac{\sigma_2}{\sigma_1}) + \frac{\sigma_1^2 + (\mu_1 - \mu_2)^2}{2\sigma_2^2} - \frac{1}{2} \right)$

$$\frac{\partial T_2}{\partial \mu_i} = \frac{(1-q_i)\mu_i}{\delta^2} + \frac{(q_i)\mu_i}{\sigma_\beta^2}, \tag{23}$$

$$\frac{\partial T_2}{\sigma_i^2} = \frac{1}{2}\left(\frac{(1-q_i)}{\delta^2} + \frac{(q_i)}{\sigma_\beta^2} - \frac{1}{\sigma_i^2}\right), \tag{24}$$

$$\begin{aligned}\frac{\partial T_2}{\partial q_i} &= -\left(\log(\frac{\delta}{\sigma_i}) + \frac{\sigma_i^2+\mu_i^2}{2\delta^2}\right) + \left(\log(\frac{\sigma_\beta}{\sigma_i}) + \frac{\sigma_i^2+\mu_i^2}{2\sigma_\beta^2}\right) \\ &= \log(\frac{\sigma_\beta}{\delta}) - \frac{\sigma_i^2+\mu_i^2}{2\delta^2} + \frac{\sigma_i^2+\mu_i^2}{2\sigma_\beta^2}. \end{aligned} \tag{25}$$

It is clear that there is just one nonzero term in $\nabla_\theta T_2$ and it can be calculated as

$$\frac{\partial T_2}{\partial \sigma_\beta^2} = \sum_{i=1}^{M} \frac{-q_i}{2\sigma_\beta^4}\left(\sigma_i^2 + \mu_i^2 - \sigma_\beta^2\right). \tag{26}$$

Hence

$$\nabla_\theta T_2 = \begin{bmatrix} 0 \\ \sum_{i=1}^{M} \frac{-q_i}{2\sigma_\beta^4}\left(\sigma_i^2+\mu_i^2-\sigma_\beta^2\right) \\ 0 \end{bmatrix}. \tag{27}$$

For the evaluation of $KL(q(u_i)||p(u_i|\theta))$ term in $T_3$, we need to apply KL divergence of two Bernoulli distributions since $q(u_i) \equiv Bern(u_i|q_i)$ and $p(u_i|\theta) \equiv Bern(\pi_1)$. Then the corresponding KL divergence can be obtained as[2]

$$KL(q(u_i)||p(u_i|\theta)) \equiv KL(Bern(u_i|q_i)||Bern(\pi_1)) = q_i \log\frac{q_i}{\pi_1} + (1-q_i)\log(\frac{1-q_i}{1-\pi_1}).$$

Then,

$$T_3 = \sum_{i=1}^{M} q_i \log\frac{q_i}{\pi_1} + (1-q_i)\log(\frac{1-q_i}{1-\pi_1}),$$

[2]Let x and y be two Bernoulli distributions with parameters $p_x$ and $p_y$. Then $KL(x||y) = p_x \log(\frac{p_x}{p_y}) + (1-p_x)\log\frac{1-p_x}{1-p_y}$.

and corresponding nonzero derivatives are:

$$\frac{\partial T_3}{\partial q_i} = \log \frac{q_i}{\pi_1} - \log \frac{1-q_i}{1-\pi_1}, \tag{28}$$

$$\frac{\partial T_3}{\partial \pi_1} = \sum_{i=1}^{M} \frac{\pi_1 - q_i}{\pi_1 - \pi_1^2}, \tag{29}$$

$$\nabla_\theta T_3 = \begin{bmatrix} \sum_{i=1}^{M} \frac{\pi_1 - q_i}{\pi_1 - \pi_1^2} \\ 0 \\ 0 \end{bmatrix}, \tag{30}$$

hence

$$\nabla_\theta \mathcal{L}_{q,\theta} = \begin{bmatrix} -\sum_{i=1}^{M} -\frac{\pi_1 - q_i}{\pi_1 - \pi_1^2} \\ -\sum_{i=1}^{M} \frac{-q_i}{2\sigma_\beta^4}\left(\sigma_i^2 + \mu_i^2 - \sigma_\beta^2\right) \\ \frac{T_A - M\sigma_0^2}{2\sigma_0^4} \end{bmatrix}. \tag{31}$$

All in all, as summarized in Table 1, we can add up the corresponding derivatives of all terms to get the derivatives of $\mathcal{L}_{q,\theta}$.

| | $T_1$ | $T_2$ | $T_3$ | $L_{q,\theta}$ |
|---|---|---|---|---|
| $\partial \mu_i$ | $\frac{1}{\sigma_0^2}\sum_{j=1}^{M}$ $a_{ij}(z_j - \sum_{k=1}^{M} a_{kj}\mu_k)$ | $\frac{(1-q_i)}{\delta^2} + \frac{(q_i)}{\sigma_\beta^2}$ | 0 | $\frac{1}{\sigma_0^2}\sum_{j=1}^{M} a_{ij}(z_j - \sum_{k=1}^{M} a_{kj}\mu_k)$ - $\frac{(1-q_i)\mu_i}{\delta^2} - \frac{(q_i)\mu_i}{\sigma_\beta^2}$ |
| $\partial \sigma_i^2$ | $\frac{-1}{4\sigma_0^2}\sum_{j=1}^{M} 2a_{ij}^2$ | $\frac{1}{2}\left(\frac{(1-q_i)\mu_i}{\delta^2} + \frac{(q_i)\mu_i}{\sigma_\beta^2} - \frac{1}{\sigma_i^2}\right)$ | 0 | $\frac{-1}{4\sigma_0^2}\sum_{j=1}^{M} 2a_{ij}^2$ - $\frac{1}{2}\left(\frac{(1-q_i)\mu_i}{\delta^2} + \frac{(q_i)\mu_i}{\sigma_\beta^2} - \frac{1}{\sigma_i^2}\right)$ |
| $\partial q_i$ | 0 | $\log(\frac{\sigma_\beta}{\delta}) - \frac{\sigma_i^2+\mu_i^2}{2\delta^2} + \frac{\sigma_i^2+\mu_i^2}{2\sigma_\beta^2}$ | $\log\frac{q_i}{\pi_1} - \log\frac{1-q_i}{1-\pi_1}$ | -$\left(\log(\frac{\sigma_\beta}{\delta}) - \frac{\sigma_i^2+\mu_i^2}{2\delta^2} + \frac{\sigma_i^2+\mu_i^2}{2\sigma_\beta^2} + \log\frac{q_i}{\pi_1} - \log\frac{1-q_i}{1-\pi_1}\right)$ |
| $\nabla_\theta$ | $\begin{bmatrix} 0 \\ 0 \\ \frac{T_A - M\sigma_0^2}{2\sigma_0^4} \end{bmatrix}$ | $\begin{bmatrix} 0 \\ \sum_{i=1}^{M} \frac{-q_i}{2\sigma_\beta^4}\left(\sigma_i^2 + \mu_i^2 - \sigma_\beta^2\right) \\ 0 \end{bmatrix}$ | $\begin{bmatrix} \sum_{i=1}^{M} \frac{\pi_1 - q_i}{\pi_1 - \pi_1^2} \\ 0 \\ 0 \end{bmatrix}$ | $\begin{bmatrix} -\sum_{i=1}^{M} \frac{\pi_1 - q_i}{\pi_1 - \pi_1^2} \\ -\sum_{i=1}^{M} \frac{-q_i}{2\sigma_\beta^4}\left(\sigma_i^2 + \mu_i^2 - \sigma_\beta^2\right) \\ \frac{T_A - M\sigma_0^2}{2\sigma_0^4} \end{bmatrix}$ |

Table 1: All partial derivatives of $\mathcal{L}_{q,\theta}$

#### Computational Complexity of Derivative Calculation

As will be seen in the following sections, we will use ADAM algorithm which requires the online calculation of derivatives at each iteration. Therefore it is important to determine the computational cost of these derivatives per iteration. To do so, we may start by examining $\frac{\partial \mathcal{L}_{q,\theta}}{\partial \mu_i}$,

$$\frac{\partial \mathcal{L}_{q,\theta}}{\partial \mu_i} = \left(\frac{1}{\sigma_0^2}\sum_{j=1}^{M} a_{ij}(z_j - \sum_{k=1}^{M} a_{kj}\mu_k)\right) - \frac{(1-q_i)\mu_i}{\delta^2} - \frac{(q_i)\mu_i}{\sigma_\beta^2}. \tag{32}$$

Note that $\frac{\partial \mathcal{L}_{q,\theta}}{\partial \mu_i}$ can be written in more compact from as

$$\frac{\partial \mathcal{L}_{q,\theta}}{\partial \boldsymbol{\mu}} = \frac{1}{\sigma_0^2}(A_1 + A_2\boldsymbol{\mu})^T - \frac{(1-\boldsymbol{q}) \odot \boldsymbol{\mu}}{\delta^2} - \frac{\boldsymbol{q} \odot \boldsymbol{\mu}}{\sigma_\beta^2},$$

where $A_1 = A\boldsymbol{z}$ and $A_2 = -AA^T$, $\odot$ is Hadamard product, $\boldsymbol{q}$ and $\boldsymbol{m}$ are the vectors that have all $q_i$ and $\mu_i$ elements, respectively. Since $A_1$ and $A_2$ can be pre-computed, the required computation per iteration is $O(M^2)$ which comes from $A_2\boldsymbol{\mu}$ term.

The derivative with respect to $\sigma_i^2$ can be written as:

$$\frac{\partial \mathcal{L}_{q,\theta}}{\partial \sigma_i^2} = \left( \frac{-1}{4\sigma_0^2} \sum_{j=1}^{M} 2a_{ij}^2 \right) - \frac{(1-q_i)}{\delta^2} - \frac{(q_i)}{\sigma_\beta^2}. \tag{33}$$

Note that the first term of $\frac{\partial \mathcal{L}_{q,\theta}}{\partial \sigma_i^2}$ is constant and can be pre-computed. Hence the required computation per iteration is $O(M)$. Regarding $\frac{\partial \mathcal{L}_{q,\theta}}{\partial q_i}$ which can be expressed as,

$$\frac{\partial \mathcal{L}_{q,\theta}}{\partial q_i} = -\left( \log(\frac{\sigma_\beta}{\delta}) - \frac{\sigma_i^2 + \mu_i^2}{2\delta^2} + \frac{\sigma_i^2 + \mu_i^2}{2\sigma_\beta^2} + \log\frac{q_i}{\pi_1} - \log\frac{1-q_i}{1-\pi_1} \right), \tag{34}$$

the computation per iteration for $\frac{\partial \mathcal{L}_{q,\theta}}{\partial q_i}$ is $O(M)$ hence total computational complexity of the algorithm per iteration is $O(M^2)$. Note that we did not mention the complexity of the calculation of $\nabla_\theta \mathcal{L}_{q,\theta}$ (which is $O(M)$). The reason is, we are not using them in ADAM algorithm hence there is no need to take them into account in this current work but planning to use them in our future work.

**Reducing Computational Complexity with Finemap-MiXeR PCA**

As mentioned above, the required computation to calculate derivatives $\frac{\partial \mathcal{L}_{q,\theta}}{\partial \mu_i}$, $\frac{\partial \mathcal{L}_{q,\theta}}{\partial \sigma_i^2}$, $\frac{\partial \mathcal{L}_{q,\theta}}{\partial_i}$ are $O(M^2)$, $O(M)$ and $O(M)$ respectively. Hence, if we can reduce the computation of $\frac{\partial \mathcal{L}_{q,\theta}}{\partial \mu_i}$ somehow, we can also reduce the required computation of the whole algorithm. Firstly let's re-write the exact formula of this derivative:

$$\frac{\partial \mathcal{L}_{q,\theta}}{\partial \boldsymbol{\mu}} = \frac{1}{\sigma_0^2}(A_1 + A_2\boldsymbol{\mu})^T - \frac{(1-\boldsymbol{q}) \odot \boldsymbol{\mu}}{\delta^2} - \frac{\boldsymbol{q} \odot \boldsymbol{\mu}}{\sigma_\beta^2}.$$

The problematic part in this expression that requires $O(M^2)$ is $A_2\boldsymbol{\mu}$ which is a MxM matrix and Mx1 vector product where $A_2 = -AA^T$. Let's recall the result of this product as

$$A_p = A_2\boldsymbol{\mu}. \tag{35}$$

Since $A_2 = -AA^T$, the columns of $A_2$ are highly correlated due to LD structure of matrix $A$. Therefore, for reducing dimensionality, we may perform Principal Component Analysis (PCA) analysis to $A_2$ to obtain its eigenvalues and corresponding eigenvectors as

$$cov(A_2) = U\Sigma U^T, \tag{36}$$

where $\Sigma$ corresponds to diagonal matrix whose diagonal elements are the sorted eigenvalues of cov($A_2$) and columns of U is the matrix whose columns are the corresponding eigenvectors. Then we can choose the first $p_c$ eigenvectors of U that covers the $pc_{thr}$=0.9999 (%99.99) of variation of $A_2$ as $A_{22} = U^T A_2$ and project $A_p$ into the new dimension as

$$A_{p2} = A_{22}\boldsymbol{\mu}, \tag{37}$$

where $A_{p2}$ is a matrix whose dimensions are $p_c$ x M. If we would like to reconstruct $A_p$ from $A_{p2}$ we can do this by multiplying $A_{p2}$ with $U$ as

$$\hat{A}_p = UA_{p2} = UU^T A_2\boldsymbol{\mu}. \tag{38}$$

Note that, since $U^T$ and $A_2$ are fixed, we may precalculate them before the iterations as $B_1 = U^T A_2$ and for each iteration the required calculations are

$$B_2 = B_1\boldsymbol{\mu}, \tag{39}$$
$$\hat{A}_p = UB_2, \tag{40}$$

where both operations above require O($p_c$ M) and since $p_c$ is mostly $p_c << M$, the required operations to compute gradients can be reduced importantly by preserving accuracy.

**Modifications for Adam Optimization**

The direct implementation of ADAM algorithm itself does not take constraints into account. On the other hand our decision variables $q_i$ and $\sigma_i^2$, by definition, have to have some constraints. In particular, $q_i$ corresponds to probability of being causal hence it

**Algorithm 1** Modified ADAM algorithm for Finemap-MiXeR

**Require:** : $\alpha$ : Stepsize
**Require:** : $\beta_1, \beta_2 \in [0, 1)$ : Exponential decay rates for the moment estimates, paper parameters are used
**Require:** : Reparametrize $q_i = \frac{1}{1+e^{-k_f o_i}}$
**Require:** : $\mathcal{L}(x, \theta)$,: Stochastic objective function with parameters $\theta$ where x=[$\mu$ $\sigma$ o]
**Require:** : $\mathrm{x}_0$ : Initial parameter vector $\mathrm{m}_0 \leftarrow 0$ (Initialize 1st moment vector)
**Require:** : $v_0 \leftarrow 0$ (Initialize 2nd moment vector) $t \leftarrow 0$ (Initialize timestep)
**while** $q_t$ not converged **do** $t \leftarrow t + 1$
$g_t \leftarrow \nabla_\theta \mathcal{L}\left(\mathrm{q}_{\mathrm{t}-1}, \theta\right)$ : (First gradients w.r.t. stochastic objective at timestep t)
$\mathrm{m_t} \leftarrow \beta_1 \cdot \mathrm{m_t} - 1 + (1 - \beta_1) \cdot \mathrm{g_t}$ (Update biased first moment estimate)
$\mathrm{V_t} \leftarrow \beta_2 \cdot \mathrm{V_t} - 1 + (1 - \beta_t) \cdot \mathrm{g_t^2}$ (Update biased second raw moment estimate)
$\hat{m}_t \leftarrow m_t / \left(1 - \beta_1^2\right)$ (Compute bias-corrected first moment estimate)
$\hat{v}_{\mathrm{t}} \leftarrow \mathrm{V_t} / \left(1 - \beta_2^2\right)$ (Compute bias-corrected second raw moment estimate)
$\mathrm{x_t} \leftarrow \mathrm{x_{t-1}} - \alpha \cdot \widehat{m}_{\mathrm{t}} / (\mathrm{eps} + \sqrt{\mathrm{vt}}\ )$ (Update parameters)
if $\sigma_i$ is smaller than 0, project into (0, $\infty$)
**end while**
**return** $\mathrm{x_t}$ (Resulting parameters)

needs to be between 0 and 1. Similarly since $\sigma_i^2$ represents variance, it needs to be non-negative. To satisfy these constraints, we may either employ Reparamtrization (REP) or Projected Gradient (PG) approaches. For optimization of $q_i$ we are using REP by reparametrizing $q_i$ with another variable $o_i$ as

$$q_i = \frac{1}{1 + e^{-k_f o_i}} \tag{41}$$

where $k_f$ is an arbitrarily chosen constant. Therefore, regardless of the optimized value of $o_i$, $q_i$ is guaranteed to be placed between 0 and 1. Here instead of optimizing with respect to $q_i$, optimization with respect to $o_i$ is performed by determining the derivative of $\mathcal{L}(q, \theta)$ with respect to $o_i$ and it can be obtain using chain rule;

$$\frac{\partial \mathcal{L}_{q,\theta}}{\partial o_i} = \frac{\partial \mathcal{L}_{q,\theta}}{\partial q_i} \frac{\partial q_i}{\partial o_i}. \tag{42}$$

For $\sigma_i^2$, we are using Projected Gradient which is basically projecting the calculated $\sigma_i^2$ to the defined space which is (0, $\infty$) in our case.

### References

[1] A. A. Shadrin, O. Frei, O. B. Smeland, F. Bettella, K. S. O'Connell, O. Gani, S. Bahrami, T. K. Uggen, S. Djurovic, D. Holland *et al.*, "Phenotype-specific differences in polygenicity and effect size distribution across functional annotation

categories revealed by ai-mixer," *Bioinformatics*, vol. 36, no. 18, pp. 4749–4756, 2020.

[2] D. P. Kingma and J. Ba, "Adam: A method for stochastic optimization," *arXiv preprint arXiv:1412.6980*, 2014.

[3] M. Titsias and M. Lázaro-Gredilla, "Doubly stochastic variational bayes for non-conjugate inference," in *International conference on machine learning*. PMLR, 2014, pp. 1971–1979.
